# Transdiagnostic variations in impulsivity and compulsivity in obsessive-compulsive disorder and gambling disorder correlate with effective connectivity in cortical-striatal-thalamic-cortical circuits

**DOI:** 10.1101/389320

**Authors:** Linden Parkes, Jeggan Tiego, Kevin Aquino, Leah Braganza, Samuel R. Chamberlain, Leonardo Fontenelle, Ben J. Harrison, Valentina Lorenzetti, Bryan Paton, Adeel Razi, Alex Fornito, Murat Yücel

## Abstract

**Background:** Individual differences in impulsivity and compulsivity is thought to underlie vulnerability to a broad range of disorders and are closely tied to cortical-striatal-thalamic-cortical (CSTC) function. However, whether impulsivity and compulsivity in clinical disorders is continuous with the healthy population and explains CSTC dysfunction across different disorders remains unclear.

**Methods:** We characterized the relationship between CSTC effective connectivity, estimated using dynamic causal modelling of functional magnetic resonance imaging data, and dimensional phenotypes of impulsivity and compulsivity in two symptomatically distinct but phenotypically related disorders, obsessive-compulsive disorder (OCD) and gambling disorder (GD). 487 online participants provided data for modelling of dimensional phenotypes. These data were combined with 34 OCD patients, 22 GD patients, and 39 healthy controls, who underwent functional magnetic resonance imaging.

**Results:** Three core dimensions were identified: disinhibition, impulsivity, and compulsivity. Patients’ scores on these dimensions were continuously distributed with the healthy participants, supporting a continuum model of psychopathology. Across all participants, higher disinhibition correlated with lower bottom-up connectivity in the dorsal circuit and increased bottom-up connectivity in the ventral circuit, and higher compulsivity correlated with reduced bottom-up connectivity in the dorsal circuit. Similar changes in effective connectivity were observed with increasing clinical severity that were not accounted for by phenotypic variation, demonstrating convergence towards behaviourally and clinically relevant changes in brain dynamics. Effective connectivity did not differ as a function of traditional diagnostic labels.

**Conclusions:** CSTC dysfunction across OCD and GD is better characterized by dimensional phenotypes than diagnostic comparisons, supporting investigation of quantitative liability phenotypes.

## Introduction

Psychiatric research is gradually shifting away from studying classically diagnosed disorders towards an understanding of the underlying constructs and mechanisms that drive maladaptive behavior (1; 2). In this context, impulsivity and compulsivity feature prominently as putative intermediate phenotypes linked to symptom variation across multiple disorders (3–7), and likely explain a substantial fraction of commonly observed comorbidities (7–9).

Historically, impulsivity and compulsivity have been quantified using scores on either behavioral tasks (e.g., response inhibition paradigms) or self-report questionnaires (10; 11), and the relation between the two constructs has been unclear (3; 5); some suggest that they are diametrically opposed on a single continuum (12; 13), whereas others propose that they are orthogonal dimensions (4; 6). Recent Confirmatory Factor Analyses (CFA) of multiple measures of impulsivity and compulsivity has shown that the constructs form two distinct but positively correlated traits, which each predict poorer quality of life (7). Using Structural Equation Modelling (SEM) of 12 self-report measures of impulsivity and compulsivity in a large normative sample, we reported evidence for a bifactor model in which a unitary, general ‘disinhibition’ dimension, characterized by high impulsivity, uncertainty intolerance and obsessive beliefs, coupled with low desire for predictability, perfectionism, and threat estimation, was the strongest predictor of the co-occurrence of addictive and obsessive-compulsive symptomatology, with residual, specific dimensions of ‘impulsivity’ and ‘compulsivity’ explaining additional unique variance (9). Thus, our model successfully captures both correlated (disinhibition) and orthogonal variance associated with different measures of impulsivity and compulsivity that are relevant to understanding behavior and psychopathology.

One implication of our model is that clinically diagnosable disorders of impulsivity and compulsivity represent extreme expressions of traits that are distributed continuously with the healthy population (the continuity hypothesis). This postulate, while consistent with the implicit assumption of the Research Domain Criteria (RDoC) initiative (1; 2), has seldom been formally tested in psychiatry. Hence, it remains unclear how subclinical variation in impulsivity and compulsivity relate to case-level psychopathology, either at the level of observable behavior or underlying neurobiology.

Here, we test the continuity hypothesis using impulsivity and compulsivity as intermediate phenotypes and diagnosed gambling disorder (GD) and obsessive-compulsive disorder (OCD) as exemplars of psychopathology at the extreme ends of these phenotypes. GD and OCD are both associated with dysfunctional levels of impulsivity and compulsivity (14–18) and have overlapping pathophysiology centered on cortical-striatal-thalamic-cortical (CSTC) circuits (19–22), which are thought to play a critical role in mediating impulsive and compulsive behaviors (5; 23–26). An advantage of studying a behavioral addiction such as GD is that it allows us to uncover pathophysiological processes without the confounding effects of substance abuse or dependence (27–29).

The CSTC circuitry of the brain comprises a series of parallel yet integrated loops that topographically connect distinct regions of frontal cortex predominantly with ipsilateral striatum and thalamus (30; 31). These circuits are functionally specialized and broadly segregate into ventral limbic, dorsal associative, and caudal sensorimotor systems (30–33). Altered functional coupling (dysconnectivity) of the dorsal and ventral circuits has been similarly implicated in both GD and OCD (19; 21; 22; 34–41), but the two disorders have not been directly compared and the degree to which any neural similarities or differences relate to variations in impulsivity and compulsivity are unclear. Some have suggested that the ventral and dorsal striatum respectively drive impulsive and compulsive behaviors, while top-down cortical projections inhibit these behaviors (5), but these assertions have not been directly tested. Moreover, most work to date has relied on simple (undirected) models of network interactions, based on correlational measures of functional coupling between regions (i.e., functional connectivity), which cannot disentangle causal top-down or bottom-up influences in CSTC circuitry.

In this study, we addressed two primary aims. First, we extended our prior modelling work (9) by combining our existing normative cohort with a new sample of healthy controls (HCs), individuals with OCD, and individuals with GD to replicate our model and formally test the continuity hypothesis; that is, that GD and OCD participants lie at the extreme ends of our quantitative phenotypes. Second, we investigated how our quantitative and clinical phenotypes relate to CSTC function. We mapped the effective connectivity (i.e., the causal interactions between brain regions) of the CSTC circuitry using Dynamic Causal Modelling (DCM) (42–45) of resting-state functional Magnetic Resonance Imaging (rs-fMRI) data and linked effective connectivity parameters to our quantitative impulsivity and compulsivity phenotypes, as well as to traditional diagnostic groupings, in a Bayesian framework. As opposed to undirected estimates of functional connectivity, our approach allowed us to distinguish top-down from bottom-up influences in CSTC circuitry, and to evaluate whether quantitative trait variation or diagnosis is a stronger correlate of brain function.

## Methods and Materials

### Participants

Participants included 487 individuals used in our previous work (9), recruited online through the Amazon Mechanical Turk community (online dataset), and 96 participants (39 HCs, 34 OCD, and 23 GD) recruited locally for neuroimaging (imaging dataset). All participants completed a self-report questionnaire battery used to model impulsivity and compulsivity. See supplement for details on recruitment and eligibility.

### Impulsivity and compulsivity

We measured diverse aspects of impulsivity and compulsivity using self-report indices that assessed the phenotypes as multidimensional constructs, had good validity and reliability, were not measures of disorder-specific severity, and were sensitive to clinical and subclinical variation. Impulsivity was measured using the 59-item UPPS-P Impulsivity scale (46; 47). Compulsivity was measured with the Obsessive Beliefs Questionnaire 44-item version (OBQ-44) and the 12-item version of the Intolerance of Uncertainty Scale (IUS-12) (48–52).

### Structural equation modelling

We modelled the dimensional structure of the self-report measures mentioned above using SEM, as per procedures described in Tiego et al (9). Briefly, item-level data from the OBQ-44, IUS-12, and UPPS-P were combined across the online and imaging datasets. Data were fit to a bifactor model that included a general disinhibition dimension as well as specific dimensions for impulsivity and compulsivity (9). Factor score estimates were obtained for the dimensional phenotypes of disinhibition, impulsivity, and compulsivity for use in subsequent analysis of the imaging dataset. Details of model estimation as well as comparison to competing models are provided in the supplement.

### Magnetic Resonance Imaging and Dynamic Causal Modelling

Functional (echoplanar imaging, EPI) and structural (T1-weighted MP-RAGE) data were acquired on a Siemens MAGNETOM Skyra 3T scanner. Scans were processed using Matlab code available online (https://github.com/lindenmp/rs-fMRI). Pre-processing and quality control was performed as per Parkes et al (53) and are detailed in the supplement. Effective connectivity was estimated using spectral DCM (spDCM) (54; 55) implemented in SPM12 r7219 (Wellcome Trust Centre for Neuroimaging, London, UK; code available at: https://github.com/lindenmp/rs-fMRI/tree/master/stats/spDCM). spDCM was developed specifically for modelling rs-fMRI data and provides a computationally efficient way to estimate effective connectivity by fitting cross spectra rather than the time series (54). Owing to projections within CSTC circuitry being predominantly ipsilateral (30; 31), we estimated CSTC effective connectivity separately in each hemisphere.

### Generation of DCM nodes

We defined functional regions of interest (ROI) that sampled key regions of the CSTC circuits implicated in impulsivity, compulsivity, GD, and OCD. The dorsal circuit comprised the dorsal striatum, anterior cingulate cortex (aCC), orbitofrontal cortex/ventromedial prefrontal cortex (OFC/vmPFC), and anterior thalamus (5; 36). The ventral circuit comprised the ventral striatum, aCC, OFC/vmPFC, dorsolateral prefrontal cortex (dlPFC), and posterior thalamus (5; 35). Dorsal and ventral striatal subregions were extracted from a striatal parcellation based on structural connectivity (32). All other ROIs were generated using a subject-specific approach based on seed-based functional connectivity from these striatal subregions, details of which can be found in the supplement. In brief, for each CSTC circuit and subject, each anatomical ROI outside the striatum was refined into a spherical DCM ROI with radius 3-mm that satisfied the following criteria: (i) within a 16-mm radius of the second-level main effect of striatal seed for the whole sample (12-mm for the thalamus); (ii) within the boundaries of the corresponding anatomical ROI; (iii) at least 20-mm away from the center-of-mass of the corresponding striatal seed; and (iv) did not overlap with DCM ROIs generated for any of the other anatomical ROIs.

### Specification and inversion of DCM at the First Level

For each participant and hemisphere, a sparse parent model (*Figure 1*) was created that modelled connectivity from the cortical DCM ROIs to the corresponding striatal DCM ROI, from the striatal DCM ROI to the corresponding thalamic DCM ROI, and from the thalamic DCM ROI back to the corresponding cortical DCM ROIs. We excluded connections within the CSTC circuits that were not relevant to our hypotheses. Hence, cortico-cortico connections were not modelled as we were primarily interested in examining the connectivity between cortex and subcortex. This procedure was repeated for each CSTC circuit and both were combined into a single DCM. To link the two circuits, the dorsal and ventral striatal subregions were connected reciprocally. Nested models were generated by systematically turning off each of the connections present in the parent model. The parent model was inverted using spDCM (**spm_dcm_fit.m**), and the nested models were inverted using Bayesian model reduction (**spm_dcm_bmr.m**) (56). Because nested models are defined only by the removal of connections included in the parent model, they differ only in terms of their priors, which allows efficient/rapid estimation of nested DCM models by using the posterior of the parent model.

**Figure 1.**
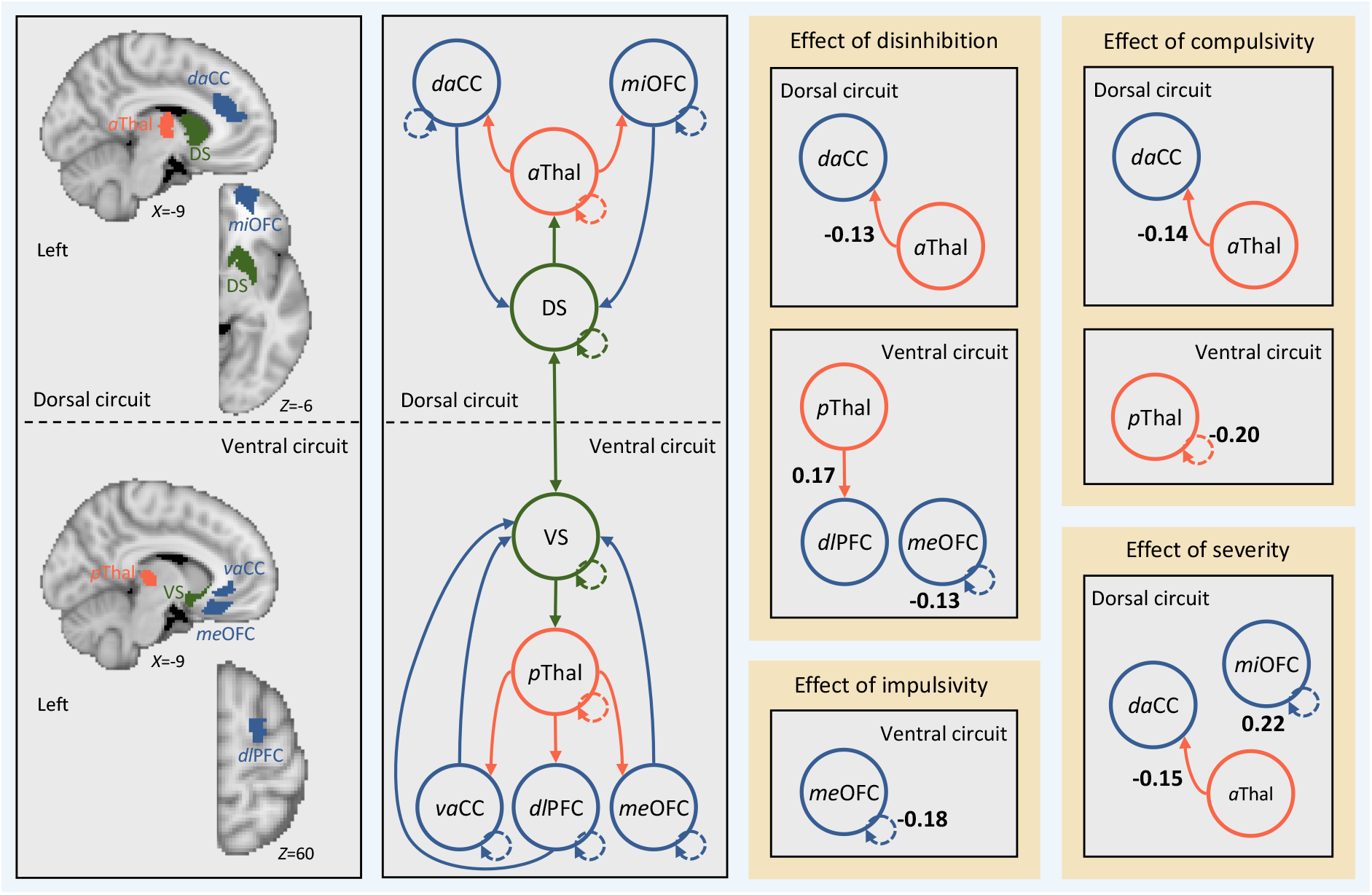
Effective connectivity between regions in cortical-striatal-thalamic-cortical (CSTC) circuits covaries with individual differences in disinhibition, compulsivity, and impulsivity. Left, brain regions in the *left* hemisphere (i.e., search volumes for subject-specific DCM ROIs) and wiring diagram representing the model that was specified. Solid lines represent connectivity between regions. Arrow heads depict direction of connection. Dashed lines represent inhibitory self-connections. Right, effects of phenotypes and clinical severity on effective connectivity within CSTC circuits in the *left* hemisphere as assessed using parametric empirical Bayes. Effective connectivity parameters are described in Hz (right panel, bold numbers), where the activity in one node influences the rate of change in the activity in another. Self-connections are (log)scaled. In both cases, the interpretaion is the same, with increasing scores on a given phenotype the effective connectivity between, or the inhibitory activity within, nodes decreases/increases as specified. The DCM ROIs have each been labeled according to which subregions of their respective anatomical ROI they were localized to during DCM ROI generation (see supplementary methods and supplementary results for details). daCC = dorsal anterior cingulate cortex, vaCC = ventral anterior cingulate cortex, miOFC = middle orbitofrontal cortex, meOFC = medial orbitofrontal cortex, aThal = anterior thalamus, pThal = posterior thalamus, dlPFC = dorsolateral prefrontal cortex, DS = dorsal striatum, VS = ventral striatum.

### Second level DCM analysis using parametric empirical Bayes

We used the parametric empirical Bayes (PEB) routines to perform second level analysis and Bayesian model averaging (**spm_dcm_peb.m** & **spm_dcm_peb_bmc.m**). PEB is a fully Bayesian and hierarchical second-level analysis framework. In hierarchical models, the posterior density over model parameters is constrained by the posterior from the level above. In group studies, this translates to modelling how within-subject effects relate to second-level group effects and differs from classical testing in that it considers the full posterior density over the parameters (i.e., both the expected connection strengths and their covariance from each first-level DCM). We used a pair of second-level PEB routines to examine: (1) the effects of disinhibition, impulsivity, compulsivity, and case-control/case-case comparisons on effective connectivity; and (2) the effects of clinical severity on effective connectivity.

For the first PEB model, a second-level design matrix space was defined using a constant in the first column (modelling the mean across the sample), the three phenotypes of disinhibition, impulsivity, and compulsivity, as well as covariates for age, gender, IQ, medication status (medicated/unmedicated), and mean framewise displacement (mFD), a summary measure of head motion (53). To examine the effect of case-control/case-case comparisons on effective connectivity, we included linear contrasts for all possible combinations of our diagnostic groups (HC, OCD, and GD). We generated the following contrasts: (i) HC>OCD; (ii) HC>GD; (iii) HC<OCD; (iv) HC<GD; (v) OCD>GD; and (vi) OCD<GD. These six contrasts were input as separate columns in the design matrix of our PEB model.

The second PEB model retained only the OCD and GD participants and included an aggregate measure of symptom severity alongside all the same variables and contrasts from the first PEB model. We indexed clinical severity separately for the OCD and GD groups using the Obsessive-Compulsive Inventory-Revised (OCI-R (57)) and the Problem Gambling Severity Index (PGSI (58)), respectively. Then, as a proxy for transdiagnostic symptom severity, we z-scored each measure within each group separately, combined across both groups to produce a single measure of severity, z-scored again, and included severity in the PEB model.

## Results

### Participants and data

579 participants were included in the phenotype modelling analyses, of which 95 participants (39 HCs, 34 OCD patients, and 22 GD patients) were from the imaging dataset. Demographics are provided in Table 1. Further exclusion was applied to the imaging dataset (see *Supplementary Results*) that yielded a final imaging dataset of 38 HC participants, 32 OCD participants, and 20 GD participants. See supplementary results for details of participant exclusion, quality control of rs-fMRI data, and co-ordinates for ROIs.

**Table 1.** Sample characteristics.

| Characteristic | HC (n=39) | OCD (n=34) | GD (n=22) | Online cohort (n=487) |
| --- | --- | --- | --- | --- |
| Age in years, mean (SD) | 34 (9.47) | 31.68 (9.40) | 36.32 (12.21) | 34.2 (9.30) |
| Male, No. (%) | 19 (49) | 16 (47) | 14 (64) | 240 (49) |
| IQ, mean (SD) | 115.31 (10.65) | 115.56 (9.09) | 111.64 (8.86) | - |
| SSRI Medication, No. (%) | 0 (0) | 20 (59) | 5 (23) | - |

*Note, online cohort is the same as used in Tiego et al (9)*

### Impulsivity and Compulsivity phenotypes

We first confirmed that the bifactor model from our prior work (9) (*Figure S1, Supplement*) provided the best fit (χ^2^(31) = 52.903, *p* = .057; RMSEA = .035; 90% *CI* = .018 −.50; CFI = .99; SRMR = .041) to this new, extended dataset, relative to several competing models (*Table S1, Supplement*). We then generated factor score estimates for each of the three phenotypes for subsequent analyses. The distributions of the factor score estimates were univariate and multivariate normal, providing evidence for a continuous distribution for the disinhibition, impulsivity, and compulsivity phenotypes that spans the non-clinical and clinical spectrum.

Next, we examined whether clinical GD and OCD participants lie at the extreme ends of our phenotypes. A MANOVA revealed significant differences between HC, OCD and GD participants (Pillai’s = .110, *F* (6,1150) = 11.164, *p* < .001; η_p_^2^ = .055, on the disinhibition (*F* (2,576) = 21.197, *p* < .001, η_p_^2^= .071), impulsivity (*F* (2,576) = 10.210, *p* < .001, η_p_^2^ = .034), and compulsivity (*F*(2,576) = 3.304, *p* = .037, η_p_^2^ = .011) phenotypes. Pairwise *post hoc* comparisons corrected using the Benjamini-Hochberg False Discovery Rate (FDR, *q* = .05) (59) revealed that the OCD (*M* = .917, *SE* = .167, 95%*CI* =.589 – 1.244, *p* < .001) and GD participants (*M* =.809, *SE* = .205, 95%*CI* =.407 – 1.211, *p* < .001) were significantly higher on disinhibition than HCs; GD participants were higher on impulsivity than HCs (*M* =.697, *SE* = .190, 95%*CI* =.352 – 1.069, *p* < .001) and OCD participants (*M* = 1.073, *SE* = .238, 95%*CI* =.605 – 1.541, *p* < .001); and OCD participants were higher on compulsivity than GD participants (*M* =.590, *SE* = .294, 95%*CI* =.101 – 1.079, *p* = .023), but not HCs (*M* = .118, *SE* = .164, 95%CI =−.204 – .441, *p* = .585).

### Effective connectivity

Having demonstrated support for the continuity hypothesis, we examined associations between scores on the disinhibition, impulsivity, and compulsivity phenotypes and CSTC effective connectivity in the imaging dataset. For all analyses, we thresholded the effective connectivity results using a posterior probability of >95%. We present effects from the left hemisphere in the main text (see *Figure S2* for right hemisphere results).

Figure 1 shows the effect of each phenotype on effective connectivity, whilst controlling for the effects of the nuisance covariates, diagnostic contrasts, and the other phenotypes. Figure 1 shows that individuals with higher scores on disinhibition exhibited (i) reduced bottom-up effective connectivity from the left anterior thalamus to the left dorsal aCC (daCC) in the dorsal circuit; (ii) increased bottom-up effective connectivity from the left posterior thalamus to the left dlPFC in the ventral circuit; and (iii) reduced inhibitory activity in the self-connection for the medial OFC (meOFC) in the ventral circuit. Individuals with higher levels of compulsivity exhibited (i) reduced bottom-up effective connectivity from the left anterior thalamus to the left daCC in the dorsal circuit; and (ii) reduced inhibitory activity in the self-connection for the posterior thalamus in the ventral circuit. Individuals with high impulsivity showed no differences in effective connectivity, but rather reduced inhibitory activity in the self-connection for the medial OFC (meOFC) in the ventral circuit. No effects were observed for any case-control or case-case contrasts. A complementary analysis using traditional frequentist analysis applied to estimates of undirected functional connectivity yielded largely null findings, demonstrating the sensitivity of our Bayesian analysis of effective connectivity (*Supplementary Results*).

Second, we examined the effect of clinical severity on effective connectivity in the OCD and GD patients. Consistent with the effects of disinhibition and compulsivity, symptom severity was associated with reduced bottom-up effective connectivity from the left anterior thalamus to the left daCC in the dorsal circuit. Critically, the effects of disinhibition and compulsivity reported above remained when controlling for the effect of severity, indicating that the effects are independent. Increased severity was also associated with increased inhibitory activity in the self-connection for the middle OFC (miOFC) in the dorsal circuit.

## Discussion

The potential benefits of understanding the neurobiology of quantitative traits that underlie risk for mental illness are widely acknowledged, and underpin the RDoC model (1; 2). Here, we first characterized the dimensional structure of impulsivity and compulsivity in a sample of non-clinical individuals and people with clinically diagnosed OCD and GD, and then examined how effective connectivity within CSTC circuitry relates to quantitative variation in these dimensional traits. Complementing our earlier work (9), we find that variance in a broad battery of impulsive and compulsive self-report measures is best explained by a bifactor model, comprising a unitary disinhibition factor with loadings from nearly all scales, coupled with specific constructs capturing residual variance in impulsivity and compulsivity. We further show that OCD and GD patients typically occupied the extreme ends of a distribution that is continuous with non-clinical individuals, providing evidence for a continuum model of psychopathology in OCD and GD. Finally, we report that high levels of disinhibition and compulsivity correlate with altered bottom-up signalling from subcortical to cortical areas in dorsal and ventral CSTC systems, and that variance in effective connectivity was better explained by quantitative, transdiagnostic variation in these constructs than traditional diagnostic categories. Together, our results support the utility of using dimensional constructs that cut across traditional diagnostic boundaries for understanding pathophysiological processes in psychiatry. Our use of DCM to distinguish between top-down and bottom-up influences in CSTC circuitry also suggests that the pathological expression of these dimensional traits may be related to altered subcortical signaling to cortical regions (4; 5).

### Disinhibition, impulsivity, compulsivity and the continuum model

We previously validated a model in a large online normative sample and found that various measures of impulsivity and compulsivity were best represented by three empirically-distinct phenotypes of disinhibition, impulsivity, and compulsivity (9). Furthermore, we demonstrated that these phenotypes explained subclinical variation in a broad range of impulsive, addictive, and obsessive-compulsive symptomatology (9). Here, we extended these findings by applying the model to an expanded sample that also included individuals with diagnosed GD and OCD. We show that a bifactor model comprising disinhibition, impulsivity, and compulsivity constructs remains the best fit to the data supporting the robustness of the previously proposed model. Furthermore, OCD and GD groups generally sat on the extreme ends of traits that had univariate and multivariate normal distributions, supporting a continuum model in which disorder represents the extreme expression of traits with a continuous population distribution. Thus, our results are in line with the basic premise of the RDoC initiative, and support characterization of the full range of variation between normal and abnormal functioning as a critical first step to developing individually targeted treatment strategies in psychiatry (2).

### Quantitative traits covary with bottom-up signalling in CSTC circuitry

Higher scores on the disinhibition dimension correlated with reduced resting-state bottom-up connectivity in the dorsal circuit and increased bottom-up connectivity in the ventral circuit. These findings demonstrate that concurrent increases in impulsivity, uncertainty intolerance, and obsessive beliefs, in conjunction with reductions in desire for predictability, perfectionism, and threat estimation are associated with divergent changes across distinct CSTC circuits. Our results are the first to demonstrate that dysfunction in these aspects of impulsivity and compulsivity may be associated with a dysfunctional behavioral drive subserved by the ventral and dorsal striatal subregions (4; 5). Previous resting-state work using functional connectivity has shown increased ventral CSTC connectivity in both OCD and GD (19; 22; 35), which is reduced in OCD via deep brain stimulation to the ventral striatum (35) and our results suggest that this effect may arise through the remediation of excessive bottom-up connectivity from the ventral striatum.

We also found that higher scores on the compulsivity-specific dimension correlated with reductions in resting-state bottom-up connectivity in the dorsal CSTC circuit. This result appears counter to previous task-based fMRI studies showing dorsal striatum hyperactivation and increased functional connectivity with the ACC in OCD patients who develop compulsive habits relative to those who do not (36). This apparent discrepancy may be related to the use of task-based versus resting-state fMRI protocols. Gillan et al (36) explain dorsal CSTC dysfunction as a deficit in goal-directed control over behavior, which leads to an over-reliance on habits. Our resting-state design did not overtly engage goal-driven behavioral systems. Hence, dysfunction in the dorsal CSTC circuit may be context-specific, such that connectivity is increased when compulsive behavior is expressed but decreased at rest. Concurrent modelling of effective connectivity during task and rest could be used to test this hypothesis in future.

Finally, effects in the dorsal circuit that overlapped with those found for disinhibition and compulsivity were also observed with increasing clinical severity, demonstrating that the variation in effective connectivity explained by our phenotypes was clinically relevant.

### The utility of effective connectivity models

Functional connectivity estimates undirected coupling between measured neurophysiological signals, whereas effective connectivity is based on a model of the causal interactions between neuronal populations that drive the measured signals (42). Previous research has shown effective connectivity is more sensitive to age-related changes than functional connectivity (60). Indeed, effective connectivity should offer a more precise characterization of pathophysiological processes that is less susceptible to various nuisance factors that can contaminate measures of functional connectivity (61). Here, we combined estimates of effective connectivity (54) with a fully Bayesian analysis framework (56). While the application of these methods in psychiatry is growing, so far they have been predominantly applied to schizophrenia (56; 62; 63). Thus, to demonstrate the utility of DCM in psychiatry, we replicated our analyses using functional connectivity and found no associations that survived correction for multiple comparisons. Our results thus support the superior sensitivity of a fully Bayesian analysis of effective connectivity for uncovering brain-behavior relationships in psychiatry.

### Limitations

Head motion is a pernicious issue in rs-fMRI data (64–66) that confounds pathophysiological inferences (53). To rigorously address this issue, we adopted state-of-the-art denoising methods (67; 68) and stringent participant exclusion criteria (53; 69) that minimized motion-related confound in our data. Our cross-sectional design is another limitation given that OCD and GD patients may express different levels of impulsivity and compulsivity throughout the course of their illness (6). Longitudinal investigations could clarify how latent phenotypes and their neural substrates change over time. More than half of our OCD patients and four of the GD patients were on SSRI medication. Covarying for medication status had no impact on our findings, but to our knowledge the precise effects of SSRIs on CSTC effective connectivity have not been investigated.

## Conclusions

Intermediate phenotypes are viewed as a promising method for understanding behavioral and biological mechanisms of risk for diverse disorders (1; 2; 70). We show that dimensional constructs related to impulsivity and compulsivity more closely track neuronal dynamics within cortico-striatal-thalamic-cortical circuits than the traditional diagnostic categories of OCD and GD. We also show that model-based estimates of effective connectivity successfully differentiate top-down and bottom-up dynamics, whereas estimates of functional connectivity yield largely null results. These findings suggest that Bayesian analysis of effective connectivity may provide a valuable tool for identifying biomarkers that cut across diagnostic boundaries.

## Supplementary Methods

### Participants

Data were obtained from two independent samples. The first consisted of 487 participants (50.7% female) aged 18 – 55 years (*M* = 34.2, *SD* = 9.3) recruited online through the Amazon Mechanical Turk community, hereafter referred to as the ‘*online dataset*’. The online dataset largely consisted of individuals from the United States (93.3%), with a small proportion from Australia (6.1%). Participants provided written informed consent prior to completing an online battery of self-report questionnaires and were reimbursed $2 (USD) per hour for their time.

The second dataset consisted of 39 HCs, 34 patients with obsessive-compulsive disorder (OCD), and 23 patients with gambling disorder (GD) that were recruited as part of a broader study, hereafter referred to as the ‘*imaging dataset*’. All participants from the imaging dataset provided informed written consent in accordance with the Monash University Human Research Ethics Committee guidelines. OCD patients were recruited from specialist clinical services located in Melbourne, Australia. GD patients and HCs were recruited from the community. To be eligible for study inclusion, all participants in the imaging dataset were required to have no lifetime history of concussion, neurological disease, or drug abuse/dependence. OCD patients were required to score >8 on the severity section of the Florida Obsessive-Compulsive Inventory (FOCI (1)) and have their diagnosis confirmed by treatment services as well as the Mini International Neuropsychiatric Interview version 5 conducted by L.P. and L.B. All GD patients engaged at least weekly in Electronic Gaming Machine (EGM) gambling, were required to score >8 on the Problem Gambling Severity Index (PGSI (2)), and had their diagnosis confirmed by the Structured Clinical Interview for DSM-IV conducted by L.P. and L.B. The presence of either depression or anxiety, indexed by the MINI, in either OCD or GD patients was not excluded so long as the OCD and GD symptoms constituted the primary cause of distress and interference in the participant’s life. Participants were excluded if they met criteria for any other psychiatric disorders, including the concurrent presence of OCD and GD.

## Measures

### Structural Equation Modelling

Data screening and preliminary analyses were conducted in IBM SPSS Statistics Version 23. Univariate outliers were identified and removed using a sequential fence procedure constructed using the upper and lower quartiles, defined as: *f*_Q_ = n/4 + (1/4), and a 2.2 multiplicative of the interquartile range (3). All confirmatory factor analysis (CFA) models were estimated in Mplus 7.2 using the covariance matrix (4). A two-step analysis strategy was used as described in Tiego et al (5): (i) item-level data obtained from the UPPS-P, IUS-12, OBQ-44, questionnaires were analyzed using separate first-order CFAs to determine their optimal latent structure; (ii) Factor score estimates representing individual differences on each of these first-order latent dimensions were entered as variables for estimation of the second-order dimensional phenotypes model. First-order CFA models of ordered categorical data were estimated using the Weighted Least Squares Means and Variance adjusted estimator (WLSMV) and Theta parameterization with item loadings and thresholds freely estimated and the error variance of latent response variables fixed at one (4; 6).

Second-order CFA of the factor score estimates generated from the first-order models were estimated using Full Information Maximum Likelihood and the Bollen-Stine Bootstrap procedure with 10,000 posterior draws (4; 7). Latent variable scaling was performed using the fixed factor method (6). *Post hoc* model fitting was performed using the Benjamini-Hochberg False Discovery Rate (FDR) and freely estimated error covariances were retained if statistically significant with the FDR set at *q* = .05. Model fit was assessed using a combination of absolute and incremental fit indices, including the chi square test statistic (χ^2^), Root Mean Square Error of Approximation (RMSEA) (ε < .05 close approximate fit; ε = .05 −.08 close approximate fit; ε = .08 – 1.0 reasonable approximate fit), Comparative Fit Index (CFI) (≥.90 = reasonable fit; ≥.95 = good fit), and Standardized Root Mean Square Residual (SRMR; <.08 = good fit), or Weighted Root Mean Residual (WRMR; ≤.950 = good fit) for categorical variables (8–12). Factor score estimates were generated in Mplus using the regression method and their reliability and validity were calculated using factor score determinacy and *H* index values (13; 14). The *H* index varies from 0 – 1, with values greater than .70 indicating adequate replicability of the factors across studies using the same variables . Factor score determinacies (ρ) also vary from 0 – 1, with values of approximately ≥.9 indicating that the factor score estimates provide a valid measure of individual differences on the corresponding latent dimensions (13; 15).

### Acquisition, pre-processing, denoising, and quality control of magnetic resonance imaging data

For the imaging dataset, a high-resolution anatomical image was obtained using a T1-weighted MP-RAGE structural scan (TE = 2.55ms, TR = 1.52s, flip angle = 9°, 208 slices with 1 mm isotropic voxels) and an eyes-closed rs-fMRI sequence was obtained using BOLD contrast sensitive gradient echoplanar imaging (EPI) (TE = 30ms, TR = 2.5s, flip angle = 90°, 189 volumes, 44 slices).

T1-weighted data were processed by removing the neck (FSL’s *robustfov*), segmenting into white matter (WM), cerebrospinal fluid (CSF), and gray matter (GM) probability maps (SPM8’s *New Segment*), and spatially warping the T1 and associated tissue maps to MNI space using the nonlinear deformation algorithm implemented in the Advanced Normalization Tools (ANTs (16)). In order to yield more specific estimates of WM and CSF signals for subsequent denoising, we applied up to five erosion cycles to the WM mask and up to two erosion cycles to the CSF mask following extraction of the ventricles and before spatial normalization (17; 18).

Prior to denoising, EPI data were processed by removing the first four volumes and applying slice-time correction (SPM8), spatial realignment (SPM8), co-registration to the T1-weighted image and nonlinear warping to MNI space using the warps derived above. The data were then linearly detrended and intensity-normalized to mode 1000 units. Then, EPI data were spatially smoothed with a 6 mm FWHM kernel, denoised using ICA-AROMA (19; 20) and regression of the mean WM/CSF signals, before being bandpass filtered (between 0.008 and 0.08 Hz) (17). Bandpass filtering was done using fast Fourier transform and suppressing frequencies outside the bandpass range.

As per recommendations in Parkes et al (17), participants were excluded from analysis if any of the following were true: (i) mean framewise displacement (mFD) was >0.2mm; (ii) FD contained >20% motion spikes, where spikes were defined as a single FD of >0.25mm; or (iii) any FDs >5mm. Additionally, we report the residual cross-subject correlation between FD and whole brain functional connectivity following denoising, quantified both as a percentage of connections significantly impacted by motion (i.e., QC-FC correlations), as well as the impact that distance between brain regions has on this effect (i.e., QC-FC distance-dependence).

### Generation of DCM nodes

The anatomical brain regions for each CSTC circuit defined in the main text cover large parts of the brain and likely contain multiple heterogenous signals, which is not ideal for connectivity analysis. As such, to generate subject-specific DCM ROIs we combined our *a priori* anatomical constraints with group-level functional neuroanatomy (21).

First, the dorsal and ventral striatal subregions were defined using a parcellation of the striatum based on structural connectivity developed in our previous work (22). Second, we mapped the seed-based functional connectivity of the dorsal and ventral striatal subregions for each subject using a whole-brain general linear model as implemented in SPM12. For each striatal subregion, subject-specific contrast images were included in a 3 x 2 factorial design (group (HC, OCD, PG) by hemisphere (left, right)). To estimate the main effect of striatal subregion at the second level, single-sample t-tests were run for each striatal subregion and each hemisphere separately, collapsing across all three groups. Third, anatomical masks were generated in each hemisphere using the following AAL region (s): (i) thalamus; (ii) aCC; (iii) OFC/vmPFC; and (iv) dlPFC. Fourth, for each striatal subregion, anatomical mask, and hemisphere, we generated a spherical search volume with a 16-mm radius (12-mm was used for the thalamus due to the smaller anatomical size of this region) centered on the maximum t-value from the second level main effect. We multiplied each search volume by the corresponding anatomical mask to ensure voxels outside our anatomical regions of interest were not included. To ensure we were not capturing effects from within the striatal subregions themselves, we also removed voxels from the search volumes that were within 20-mm of the center of mass of the corresponding striatal subregion and subtracted voxels that overlapped with other search volumes (this only occurred for the aCC for the ventral CSTC circuit). This resulted in a set of search volumes for each CSTC circuit that captured the group level functional connectivity between each striatal subregion and the corresponding anatomical masks. Finally, for each subject and search volume, we found the maximal functional connectivity value (using the first-level contrast images) and generated a sphere with radius 3-mm as the DCM ROI. For each DCM ROI, time series were extracted as the principal eigenvariate of all voxel time series.

In summary, the above procedure resulted in subject-specific DCM ROIs with radius 3-mm that satisfied the following criteria: (i) within a 16-mm radius of the second-level main effect of striatal seed for the whole sample (12-mm for the thalamus); (ii) within the boundaries of the corresponding anatomical ROI; (iii) at least 20-mm away from the center-of-mass of the corresponding striatal seed; and (iv) did not overlap with DCM ROIs generated for any of the other anatomical ROIs.

## Supplementary Results

### Participant exclusion

Of the 487 participants from the online dataset, three were excluded because of outlying scores on the phenotypes from our bifactor model (see below). Of the 96 participants from the imaging dataset, one individual from the GD group was excluded due to outlying scores on phenotypes from our bifactor model (see below). This yielded a final phenotype modelling sample of 579 participants, of which a subset of 39 HC participants, 34 OCD participants, and 22 GD participants underwent imaging. Following phenotype modelling, four more individuals from the imaging dataset were excluded due to excessive motion (one from the HC group, one from OCD, and two from GD) and one individual from the OCD group was excluded due to poor EPI quality. This yielded a final imaging sample of 38 HC participants, 32 OCD participants, and 20 GD participants.

### Structural Equation Modelling

Results from the first-order CFA revealed that bifactor models provided a reasonable fit to the data for the UPPS-P (χ^2^ (1589) = 2854.092, *p* <.001; RMSEA = .037; 90%CI = .035 −.039; CFI = .976; WRMR = 1.346), IUS-12 (χ^2^ (31) = 43.464, *p* = .068; RMSEA = .026; 90%CI = .000 −.043; CFI = .999; WRMR = .314), OBQ-44 (χ^2^ (355) = 3931.902, *p* <.001; RMSEA = .079; 90%CI = .076 −.081; CFI = .916; WRMR = 1.727) in the combined sample, replicating previous results reported for the American Mechanical Turk sample (5). Factor score estimates were generated for each of these 12 latent dimensions and screened for univariate outliers (UPPS-P Impulsivity General = 2; OBQ-44 Obsessive Beliefs General = 1; IUS-12 Importance & Control of Thoughts - Specific = 1; IUS-12 Responsibility & Threat Estimation – Specific = 1). The correlations matrix for the factor score estimates used in the second-order CFA models are displayed in Table S1. The Overlapping Dimensional Phenotypes bifactor model (see Figure S1) was replicated from the previous study (5) and provided an acceptable fit to the data (χ^2^ (31) = 52.903, *p* = .057; RMSEA = .035; 90%CI = .018 −.050;CFI = .990; SRMR = .041) with 12 error covariances freely estimated (FDR *q* < .05). Several competing models were also estimated to determine if they provided a better fit, including one-, two-, and three-factor models. Fit statistics for these competing models are provided in Table S2. Similar to results reported in Tiego et al (5), none of these competing models provided an acceptable fit to the data. Furthermore, several variables did not load significantly on their respective factors and had to be dropped from the One Factor (Intolerance of Uncertainty – General), Two-Factor A (RT-, ICT-, & Need for Predictability in the Face of Uncertainty-Specific); Two-Factor B and Three-Factor (IUS-12 & OBQ-44 General) models, suggesting that they did not adequately capture the covariances in the data.

From the bifactor model, construct replicability was acceptable for disinhibition (.779), impulsivity (.771), and compulsivity (.721), suggesting these factors would be reliably estimated across studies using the same variables (14; 23). Factor determinancies were high for disinhibition (ρ = .978), impulsivity (ρ = .903), and compulsivity (ρ = .930), indicating that the factor scores estmates provided accurate measurements of the underlying latent dimensions (13). Intercorrelations between the factor scores estimates were weak (disinhibition with impulsivity *r* = .078, *p* = .058 & compulsivity *r* = −.100, *p* = .016; impulsivity with compulsivity *r* = −.152, *p* < .001), demonstrating that the factor score estimates for each phenotype were not contaminated by variance from the other two factors (13). Factor score estimates were generated for the disinhibition, impulsivity, and compulsivity dimensions as measured in the final Overlapping Dimensional Phenotypes bifactor model (Figure S1). The distributions of factor score estimates were screened for univariate outliers, with one GD participant exhibiting Impulsivity exceeding 2.2 times the interquartile range, and results of three HC participants from the online dataset exceeding the critical Malhalanobis distance (χ^2^ (3) >16.266, *p* < .001) (3; 24). As mentioned above, these participants were excluded from further analyses. The distributions of the factor score estimates were univariate and multivariate normal based on the skewness and kurtosis statistic divided by their standard errors for disinhibition (*Z =* 1.257, *p* = .209; *Z* = −.340, *p* = .725), impulsivity (Z = 1.794, *p* = .073; *Z* = 1.223, *p* = .220), and compulsivity (Z = 1.554, *p* = .120; *Z* = −1.202, *p* = .230) and as evaluated by Small’s test of multivariate skewness (χ^2^ (3) = 7.014, *p* = .072) and kurtosis (χ^2^ (3) = 3.207, *p* = .361) (24–26). This suggests that values for these latent variables obtained in the OCD and GD groups were continuous with the non-clinical population, consistent with the assumption of normally distributed dimensional phenotypes.

**Table S1.** Intercorrelations Amongst the First-Order Factor Score Estimates Used as Variables for the Second-Order Confirmatory Factor Analysis Models

|  | 1 | 2 | 3 | 4 | 5 | 6 | 7 | 8 | 9 | 10 | 11 | 12 |
| --- | --- | --- | --- | --- | --- | --- | --- | --- | --- | --- | --- | --- |
| 1. Age |  |  |  |  |  |  |  |  |  |  |  |  |
| 2. Impulsivity | -.018 |  |  |  |  |  |  |  |  |  |  |  |
| 3. Urgency | -.016 | .131** |  |  |  |  |  |  |  |  |  |  |
| 4. Sensation Seek | -.123** | .085* | .596*** |  |  |  |  |  |  |  |  |  |
| 5. Premeditation | .016 | .085* | .429*** | .421*** |  |  |  |  |  |  |  |  |
| 6. Perseverance | -.032 | .057 | .214*** | .051 | .458*** |  |  |  |  |  |  |  |
| 7. Intolerance | -.037 | .394*** | -.055 | -.235*** | -.424*** | -.141** |  |  |  |  |  |  |
| 8. Paralysis | -.101* | .372*** | .185*** | .065 | .121** | .335*** | .097* |  |  |  |  |  |
| 9. Predictability | .035 | -.378*** | -.211*** | -.104* | -.308 | -.378*** | .087* | -.712*** |  |  |  |  |
| 10. Obsessive | -.086* | .394*** | .117** | -.060 | -.241*** | -.078 | .616*** | .171*** | -.039 |  |  |  |
| 11. Control | -.040 | .188*** | .280*** | .064 | .217*** | .100* | -.012 | .210*** | -.257*** | .082* |  |  |
| 12. Perfection | .021 | -.169*** | -.159*** | -.101* | -.287*** | -.328*** | .176*** | -.142*** | .255*** | .066 | -.113** |  |
| 13. Threat | .097* | -.315*** | -.172*** | -.006 | -.129** | -.255*** | -.053 | -.387*** | .413*** | .016 | -.324*** | .141** |

Note. Impulsivity = Impulsivity-General (UPPS-P); Urgency = Negative and Positive Urgency-Specific (UPPS-P); Sensation Seek = Sensation Seeking-Specific (UPPS-P); Premeditation = Lack of Premeditation Specific (UPPS-P); Perseverance = Lack of Perseverance-Specific (UPPS-P); Intolerance = Intolerance of Uncertainty-General (IUS-12); Paralysis = Paralysis of Cognition and Action in the Face of Uncertainty-Specific (IUS-12); Predictability = Desire for Predictability-Specific (IUS-12); Obsessive = Obsessive Beliefs General (OBQ-44); Perfectionism = Perfectionism Specific (OBQ-44); Importance = Importance and Control of Thoughts Specific (OBQ-44); Responsibility = Responsibility and Threat Assessment (OBQ-44). \*\*\* $p < .001$ ; \*\* $p < .01$ ; \* $p < .05$ . $N = 583$ .

**Table S2.** Summary of Fit Statistics for the Competing Second-Order Confirmatory Factor Analysis Models of Impulsivity and Compulsivity

|  | **Model** | **df** | **$\chi^2$** | ***p*** | **RMSEA (90%CI)** | **SRMR** | **CFI** | **AIC** |
| --- | --- | --- | --- | --- | --- | --- | --- | --- |
| 1 | Three Factor <sup>1,2</sup>,<br>(UPPS-P, OBQ-44,<br>IUS-12) | 30 | 302.836 | <.001 | .125 (.112 - .138) | .079 | .829 | 12381.555 |
| 2 | Two Factor A <sup>1,3</sup><br>(Impulsivity &<br>Compulsivity) | 24 | 526.127 | <.001 | .189 (.176 - .204) | .139 | .656 | 12839.687 |
| 3 | Two Factor B <sup>1,2</sup><br>(separate<br>Intolerance to<br>Uncertainty<br>factor) | 24 | 62.417 | .001 | .052 (.037 - .069) | .035 | .976 | 12153.136 |
| 4 | One Factor <sup>1,4</sup> | 34 | 247.257 | <.001 | .104 (.092 - .116) | .064 | .883 | 13758.133 |
| 5 | Bifactor <sup>1</sup><br>(Overlapping<br>Dimensional<br>Phenotypes) | 31 | 52.903 | .057 | .035 (.018 - .050) | .041 | .990 | 14721.144 |

*Note.* *df* = Degrees of Freedom; $\chi^2$ = Chi square value for test of model fit estimated using Maximum Likelihood; *p* = significance value of the chi square test statistic using the Bollen-Stine Bootstrap with 10,000 posterior draws; RMSEA = Root Mean Square Error of Approximation; *CI* = Confidence Interval; SRMR = Standardized Root Mean Residual; CFI = Comparative Fit Index; AIC = Akaike Information Criterion. <sup>1</sup>Included freely estimated error covariances; <sup>2</sup>No IUS-12 or OBQ-44 general variables; <sup>3</sup>No RT-, ICT-, & Need for Predictability in the Face of Uncertainty-Specific variables; <sup>4</sup>No IUS-12 General variable.

**Figure S1.**
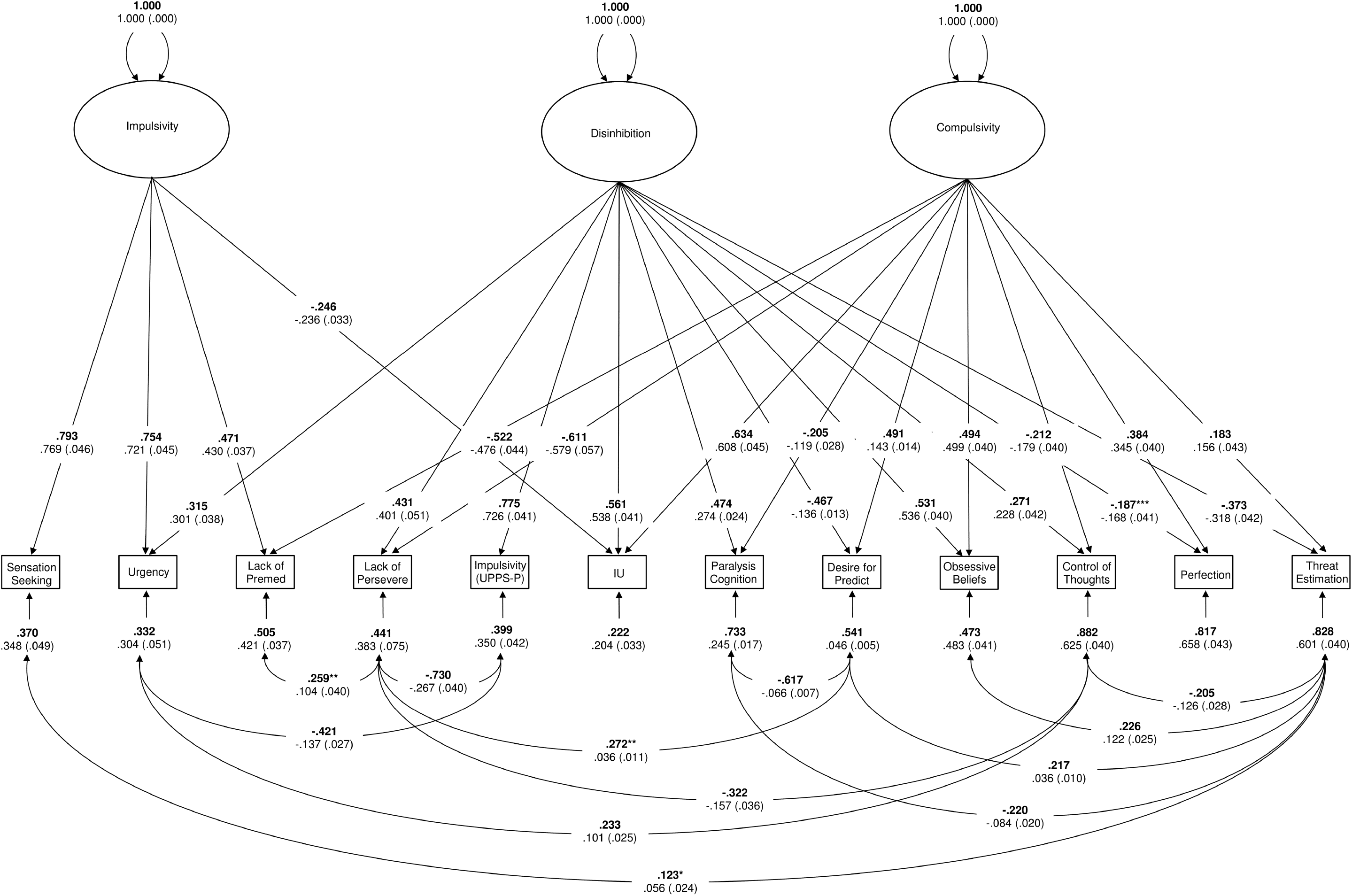
Overlapping Dimensional Phenotypes (bifactor) model, consisting of a general Disinhibition factor, and two group factors, Impulsivity and Compulsivity.

### Quality control of rs-fMRI scans

Adequate control over motion-related artefacts in rs-fMRI is crucial to conducting group analyses (17; 27–29). The mFD averaged over the clinical dataset was 0.07±0.03, indicating the level of motion was low following exclusion. At the group level, mFD was the same across HC (mFD = 0.07±0.03), OCD (mFD = 0.07±0.03), and GD (mFD = 0.07±0.03) groups. Only 3.73% of whole brain functional connections were significantly correlated (*p* <.05, uncorrected; absolute median QC-FC correlation was 0.10) with motion following denoising with ICA-AROMA and the distance-dependence on this effect was 0.04 (Spearman’s ρ), indicating that motion contamination was low in our data.

### DCM ROI generation

As mentioned above, we generated participant-specific DCM ROIs that were within a 16-mm (12-mm for the thalamus) radius of the second-level main effect of seed for each anatomical ROI in our model. We found that the dorsal and ventral striatal subregions functionally connected to distinct subregions of our anatomical ROIs, these foci are listed below in Table S3. Specifically, we found that the dorsal striatum preferentially connected to the dorsal aCC (daCC), the middle OFC (miOFC) and the anterior thalamus, whereas the ventral striatum preferentially connected to the ventral aCC (vaCC), the medial OFC (meOFC), and the posterior thalamus. The second-level functional foci for each of these regions were the center points that constrained the generation of our participant-specific DCM ROIs for the dorsal and ventral CSTC circuits.

**Table S3.** MNI co-ordinates of peak group-level connectivity within cortico-striatal-thalamic-cortical circuits. These co-ordinates formed the centre points of search volumes used to define participant-specific ROIs for dynamic causal modelling.

|  | Dorsal CSTC circuit (x,y,z) |  | Ventral CSTC circuit (x,y,z) |  |
| --- | --- | --- | --- | --- |
|  | left | right | left | right |
| Thalamus | -8,-4,8 | 2,-4,10 | -16,-28,2 | 12,-28,4 |
| aCC | -10,40,26 | 10,40,6 | -8,24,-4 | 4,28,-6 |
| OFC | -22,56,2 | 22,62,0 | -12,24,-8 | 4,30,-12 |
| dIPFC | N/A | N/A | -24,-4,74 | 50,-8,56 |

### Effective connectivity

In the main text we presented results in the left hemisphere, here we present results for the right hemisphere. Results are shown below in Figure S2. Similar to the left hemisphere, we found that none of our diagnostic group contrasts yielded any supra-threshold effect on effective connectivity between regions in the CSTC circuits. However, unlike the left hemisphere, we found that disinhibition, compulsivity, and impulsivity also had little impact on effective connectivity between CSTC regions, suggesting our results were largely lateralized to the left hemisphere.

**Figure S2.**
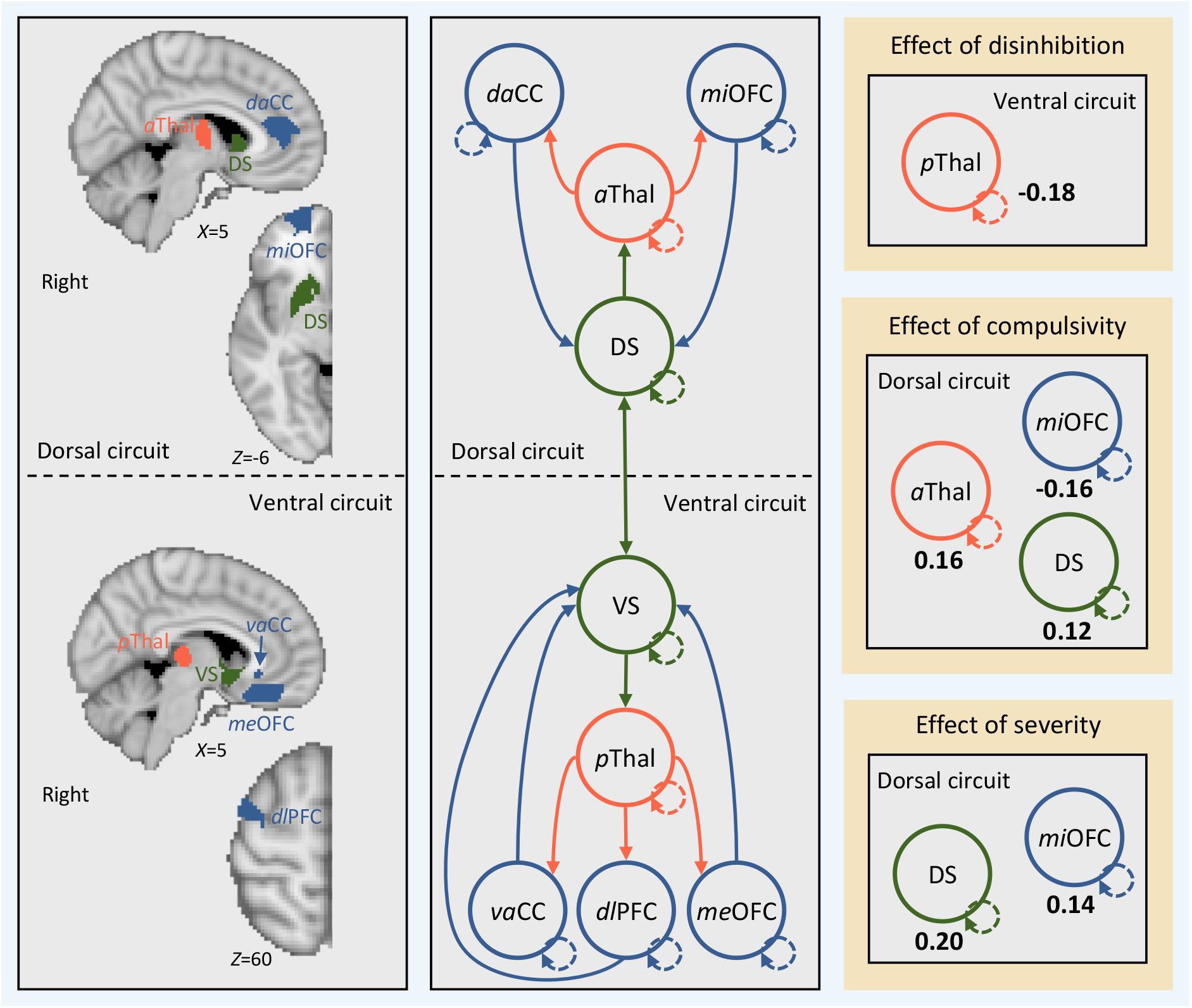
Effective connectivity between regions in cortical-striatal-thalamic-cortical (CSTC) circuits in the *right* hemisphere covaries with individual differences in disinhibition, compulsivity, and impulsivity. Left, brain regions (i.e., search volumes for subject-specific DCM ROIs) and wiring diagram representing the model that was specified. Solid lines represent connectivity between regions. Arrow heads depict direction of connection. Dashed lines represent inhibitory self-connections. Right, effects of phenotypes and clinical severity on effective connectivity within CSTC circuits in the *right* hemisphere as assessed using parametric empirical Bayes. Effective connectivity parameters are described in Hz (right panel, bold numbers), where the activity in one node influences the rate of change in the activity in another. Self-connections are (log)scaled. In both cases, the interpretaion is the same, with increasing scores on a given phenotype the effective connectivity between, or the inhibitory activity within, nodes decreases/increases as specified. The DCM ROIs have each been labeled according to which subregions of their respective anatomical ROI they were localized to during DCM ROI generation (see supplementary methods and supplementary results for details). daCC = dorsal anterior cingulate cortex, vaCC = ventral anterior cingulate cortex, miOFC = middle orbitofrontal cortex, meOFC = medial orbitofrontal cortex, aThal = anterior thalamus, pThal = posterior thalamus, dlPFC = dorsolateral prefrontal cortex, DS = dorsal striatum, VS = ventral striatum.

### Analysis using functional connectivity

In the main text we present results of a fully Bayesian analysis of effective connectivity and show that phenotypes from our bifactor model (5) covary with bottom-up connectivity in CSTC circuits. Here, we contrast this Bayesian analysis of effective connectivity with frequentist analysis of functional connectivity. We estimated functional connectivity as a Pearson correlation between the eigenvariate time series from the DCM ROIs that elicited an effect in our PEB analysis (see main text), not including the self-connections. This resulted in undirected functional connectivity being estimated between the anterior thalamus and daCC from the dorsal CSTC circuit as well as between the posterior thalamus and dlPFC from the ventral CSTC circuit. The relationship between functional connectivity and the phenotypes was assessed using a general linear model that included the same nuisance covariates as in the DCM analysis (i.e., age, gender, IQ, medication status, and mean FD). None of the model coefficients were significant at *p*<0.05, corrected for multiple comparisons. The negative relationship between compulsivity and anterior thalamus-daCC functional connectivity in the dorsal CSTC circuit was marginally significant at uncorrected levels (*r* = −0.05, *p* = 0.049). These results demonstrate that estimates of effective connectivity derived from DCM analyzed using Bayesian methods are more sensitive to relationships with our dimensional phenotypes.

## Acknowledgements

L.P. was supported by an Australian Postgraduate Award. J.T. was supported by National Health and Medical Research Council (ID:1002458, 1046054). A.F. was supported by the Charles and Sylvia Viertel Foundation, the Australian Research Council (ID: FT130100589) and the National Health and Medical Research Council (ID: 3251213, 3251250, 3251392). M.Y. was supported by a National Health and Medical Research Council Fellowship (ID: 1117188), Monash University and the David Winston Turner Endowment Fund.

## References

1. Insel T, Cuthbert B, Garvey M, Heinssen R, Pine DS, Quinn K, et al. (2010): Research Domain Criteria (RDoC): Toward a New Classification Framework for Research on Mental Disorders. AJP. 167: 748–751.

2. Cuthbert BN, Insel TR (2013): Toward the future of psychiatric diagnosis: the seven pillars of RDoC. BMC Med. 11: 1–8.

3. Dalley JW, Everitt BJ, Robbins TW (2011): Impulsivity, Compulsivity, and Top-Down Cognitive Control. Neuron. 69: 680–694.

4. Fineberg NA, Potenza MN, Chamberlain SR, Berlin HA, Menzies L, Bechara A, et al. (2010): Probing Compulsive and Impulsive Behaviors, from Animal Models to Endophenotypes: A Narrative Review. Neuropsychopharmacology. 35: 591–604.

5. Fineberg NA, Chamberlain SR, Goudriaan AE, Stein DJ, Vanderschuren LJMJ, Gillan CM, et al. (2014): New developments in human neurocognition: clinical, genetic, and brain imaging correlates of impulsivity and compulsivity. CNS Spectr. 19: 69–89.

6. Fontenelle LF, Oostermeijer S, Harrison BJ, Pantelis C, Y cel M (2011): Obsessive-Compulsive Disorder, Impulse Control Disorders and Drug Addiction. Drugs. 71: 827–840.

7. Chamberlain SR, Stochl J, Redden SA, Grant JE (2017): Latent traits of impulsivity and compulsivity: toward dimensional psychiatry. Psychol Med. 13: 1–12.

8. Gillan CM, Kosinski M, Whelan R, Phelps EA, Daw ND (2016): Characterizing a psychiatric symptom dimension related to deficits in goal-directed control. eLife. 5: e94778–24.

9. Tiego J, Oostermeijer S, Prochazkova L, Parkes L, Dawson A, Youssef G, et al. (2018): Overlapping Dimensional Phenotypes of Impulsivity and Compulsivity. Retrieved from https://osf.io/qa9eh/.

10. Robbins TW, Gillan CM, Smith DG, de Wit S, Ersche KD (2012): Neurocognitive endophenotypes of impulsivity and compulsivity: towards dimensional psychiatry. Trends in Cognitive Sciences. 16: 81–91.

11. Cyders MA, Coskunpinar A (2011): Measurement of constructs using self-report and behavioral lab tasks: Is there overlap in nomothetic span and construct representation for impulsivity? Clinical Psychology Review. 31: 965–982.

12. Hollander E (1993): Obsessive-Compulsive Spectrum Disorders: An Overview. Psychiatric Annals. 23: 355–358.

13. Hollander E, Benzaquen SD (1997): The obsessive-compulsive spectrum disorders. International Review of Psychiatry. 9: 99–110.

14. Grassi G, Pallanti S, Righi L, Figee M, Mantione M, Denys D, et al. (2015): Think twice: Impulsivity and decision making in obsessive-compulsive disorder. Journal of Behavioral Addictions. 4: 263–272.

15. Tavares H, Gentil V (2007): Pathological gambling and obsessive-compulsive disorder: towards a spectrum of disorders of volition. Rev Bras Psiquiatr, 2nd ed. 29: 107–117.

16. Prochazkova L, Parkes L, Dawson A, Youssef G, Ferreira GM, Lorenzetti V, et al. (2017): Unpacking the role of self-reported compulsivity and impulsivity in obsessive-compulsive disorder. CNS Spectr. 21: 1–8.

17. Verdejo-García A, Lawrence AJ, Clark L (2008): Impulsivity as a vulnerability marker for substance-use disorders: Review of findings from high-risk research, problem gamblers and genetic association studies. Neuroscience and Biobehavioral Reviews. 32: 777–810.

18. van Timmeren T, Daams JG, van Holst RJ, Goudriaan AE (2018): Compulsivity-related neurocognitive performance deficits in gambling disorder: A systematic review and meta-analysis. Neuroscience and Biobehavioral Reviews. 84: 204–217.

19. Ben J Harrison, Pujol J, Cardoner N, Deus J, Alonso P, López-Solà M, et al. (2013): Brain Corticostriatal Systems and the Major Clinical Symptom Dimensions of Obsessive-Compulsive Disorder. BPS. 73: 321–328.

20. Jung WH, Yücel M, Yun J-Y, Yoon YB, Cho KIK, Parkes L, et al. (2016): Altered functional network architecture in orbitofronto-striato-thalamic circuit of unmedicated patients with obsessive-compulsive disorder. Hum Brain Mapp. 38: 109–119.

21. Peters J, Miedl SF, Büchel C (2013): Elevated Functional Connectivity in a Striatal-Amygdala Circuit in Pathological Gamblers. (S. Gilbert, editor) PLoS ONE. 8: e74353–7.

22. Koehler S, Ovadia-Caro S, van der Meer E, Villringer A, Heinz A, Romanczuk-Seiferth N, Margulies DS (2013): Increased Functional Connectivity between Prefrontal Cortex and Reward System in Pathological Gambling. (Y.-F. Zang, editor) PLoS ONE. 8: e84565–13.

23. Everitt BJ, Robbins TW (2013): From the ventral to the dorsal striatum: Devolving views of their roles in drug addiction. Neuroscience and Biobehavioral Reviews. 1–9.

24. Everitt BJ, Robbins TW (2005): Neural systems of reinforcement for drug addiction: from actions to habits to compulsion. Nat Neurosci. 8: 1481–1489.

25. Everitt BJ, Belin D, Economidou D, Pelloux Y, Dalley JW, Robbins TW (2008): Neural mechanisms underlying the vulnerability to develop compulsive drug-seeking habits and addiction. Philosophical Transactions of the Royal Society B: Biological Sciences. 363: 3125–3135.

26. Gillan CM, Robbins TW, Sahakian BJ, van den Heuvel OA, van Wingen G (2016): The role of habit in compulsivity. European Neuropsychopharmacology. 1–13.

27. Goudriaan AE, Oosterlaan J, de Beurs E, van den Brink W (2004): Pathological gambling: a comprehensive review of biobehavioral findings. Neuroscience and Biobehavioral Reviews. 28: 123–141.

28. Clark L, Limbrick-Oldfield EH (2013): Disordered gambling: a behavioral addiction. Current Opinion in Neurobiology. 23: 655–659.

29. Limbrick-Oldfield EH, van Holst RJ, Clark L (2013): Fronto-striatal dysregulation in drug addiction and pathological gambling: Consistent inconsistencies? YNICL. 2: 385–393.

30. Haber SN (2003): The primate basal ganglia: parallel and integrative networks. Journal of Chemical Neuroanatomy. 26: 317–330.

31. Haber SN, Knutson B (2009): The Reward Circuit: Linking Primate Anatomy and Human Imaging. Neuropsychopharmacology. 35: 4–26.

32. Parkes L, Fulcher BD, Yücel M, Fornito A (2017): Transcriptional signatures of connectomic subregions of the human striatum. Genes, Brain and Behavior. 25: 1176–17.

33. Anderson KM, Krienen FM, Choi EY, Reinen JM, Yeo BTT, Holmes AJ (2018): Gene expression links functional networks across cortex and striatum. Nature Communications. 9: 1–14.

34. Harrison BJ, Soriano-Mas C, Pujol J, Ortiz H, L pez-Sol M, Hern ndez-Ribas R, et al. (2009): Altered Corticostriatal Functional Connectivity in Obsessive-compulsive Disorder. Arch Gen Psychiatry. 66: 1189–12.

35. Figee M, Luigjes J, Smolders R, Valencia-Alfonso C-E, van Wingen G, de Kwaasteniet B, et al. (2013): Deep brain stimulation restores frontostriatal network activity in obsessive-compulsive disorder. Nat Neurosci. 16: 386–387.

36. Gillan CM, Apergis-Schoute AM, Morein-Zamir S, Urcelay GP, Sule A, Fineberg NA, et al. (2015): Functional Neuroimaging of Avoidance Habits in Obsessive-Compulsive Disorder. AJP. 172: 284–293.

37. Reuter J, Raedler T, Rose M, Hand I, Gläscher J, Büchel C (2005): Pathological gambling is linked to reduced activation of the mesolimbic reward system. Nat Neurosci. 8: 147–148.

38. van Holst RJ, Veltman DJ, Büchel C, van den Brink W, Goudriaan AE (2012): Distorted Expectancy Coding in Problem Gambling: Is the Addictive in the Anticipation? BPS. 71: 741–748.

39. Balodis IM, Kober H, Worhunsky PD, Stevens MC, Pearlson GD, Potenza MN (2012): Diminished Frontostriatal Activity During Processing of Monetary Rewards and Losses in Pathological Gambling. BPS. 71: 749–757.

40. Choi J-S, Shin Y-C, Jung WH, Jang JH, Kang D-H, Choi C-H, et al. (2012): Altered Brain Activity during Reward Anticipation in Pathological Gambling and Obsessive-Compulsive Disorder. (L. Fontenelle, editor) PLoS ONE. 7: e45938–8.

41. Worhunsky PD, Malison RT, Rogers RD, Potenza MN (2014): Altered neural correlates of reward and loss processing during simulated slot-machine fMRI in pathological gambling and cocaine dependence. Drug and Alcohol Dependence. 145: 77–86.

42. Friston KJ, Harrison L, Penny W (2003): Dynamic causal modelling. NeuroImage. 19: 1273–1302.

43. Stephan KE, Penny WD, Moran RJ, Ouden den HEM, Daunizeau J, Friston KJ (2010): Ten simple rules for dynamic causal modeling. NeuroImage. 49: 3099–3109.

44. Razi A, Friston KJ (2016): The Connected Brain: Causality, models, and intrinsic dynamics. IEEE Signal Process Mag. 33: 14–35.

45. Razi A, Seghier ML, Zhou Y, McColgan P, Zeidman P, Park H-J, et al. (2017): Large-scale DCMs for resting state fMRI. 1–20.

46. Cyders MA, Smith GT, Spillane NS, Fischer S, Annus AM, Peterson C (2007): Integration of impulsivity and positive mood to predict risky behavior: Development and validation of a measure of positive urgency. Psychological Assessment. 19: 107–118.

47. Whiteside SP, Lynam DR (2001): The Five Factor Model and impulsivity: using a structural model of personality to understand impulsivity. Personality and Individual Differences. 30: 669–689.

48. Birrell J, Meares K, Wilkinson A, Freeston M (2011): Toward a definition of intolerance of uncertainty: A review of factor analytical studies of the Intolerance of Uncertainty Scale. Clinical Psychology Review. 31: 1198–1208.

49. Carleton RN, Norton MAPJ, Asmundson GJG (2007): Fearing the unknown: A short version of the Intolerance of Uncertainty Scale. Journal of Anxiety Disorders. 21: 105–117.

50. Gentes EL, Ruscio AM (2011): A meta-analysis of the relation of intolerance of uncertainty to symptoms of generalized anxiety disorder, major depressive disorder, and obsessive–compulsive disorder. Clinical Psychology Review. 31: 923–933.

51. Obsessive Compulsive Cognitions Working Group (2005): Psychometric validation of the obsessive belief questionnaire and interpretation of intrusions inventory—Part 2: Factor analyses and testing of a brief version. Behaviour Research and Therapy. 43: 1527–1542.

52. Myers SG, Fisher PL, Wells A (2008): Belief domains of the Obsessive Beliefs Questionnaire-44 (OBQ-44) and their specific relationship with obsessive–compulsive symptoms. Journal of Anxiety Disorders. 22: 475–484.

53. Parkes L, Fulcher B, Yücel M, Fornito A (2018): An evaluation of the efficacy, reliability, and sensitivity of motion correction strategies for resting-state functional MRI. NeuroImage. 171: 415–436.

54. Friston KJ, Kahan J, Biswal B, Razi A (2014): A DCM for resting state fMRI. NeuroImage. 94: 396–407.

55. Razi A, Kahan J, Rees G, Friston KJ (2015): Construct validation of a DCM for resting state fMRI. NeuroImage. 106: 1–14.

56. Friston KJ, Litvak V, Oswal A, Razi A, Stephan KE, van Wijk BCM, et al. (2016): Bayesian model reduction and empirical Bayes for group (DCM) studies. NeuroImage. 128: 413–431.

57. Foa EB, Huppert JD, Leiberg S, Langner R, Kichic R, Hajcak G, Salkovskis PM (2002): The Obsessive-Complusive Inventory: Development and validation of a short version. Psychological Assessment. 14: 485–495.

58. Ferris JA, Wynne HJ (2001): The Canadian problem gambling index. Ottawa, ON.

59. Benjamini Y, Hochberg Y (1995): Controlling the False Discovery Rate: A Practical and Powerful Approach to Multiple Testing. Journal of the Royal Statistical Society Series B (Methodological). 57: 289–300.

60. Tsvetanov KA, Henson RNA, Tyler LK, Razi A, Geerligs L, Ham TE, et al. (2016): Extrinsic and Intrinsic Brain Network Connectivity Maintains Cognition across the Lifespan Despite Accelerated Decay of Regional Brain Activation. Journal of Neuroscience. 36: 3115–3126.

61. Friston KJ (2011): Functional and Effective Connectivity: A Review. Brain Connectivity. 1: 13–36.

62. Stephan KE, Iglesias S, Heinzle J, Diaconescu AO (2015): Translational Perspectives for Computational Neuroimaging. Neuron. 87: 716–732.

63. Zhou Y, Zeidman P, Wu S, Razi A, Chen C, Yang L, et al. (2018): Altered intrinsic and extrinsic connectivity in schizophrenia. YNICL. 17: 704–716.

64. Power JD, Schlaggar BL, Petersen SE (2015): Recent progress and outstanding issues in motion correction in resting state fMRI. NeuroImage. 105: 536–551.

65. Satterthwaite TD, Wolf DH, Loughead J, Ruparel K, Elliott MA, Hakonarson H, et al. (2012): Impact of in-scanner head motion on multiple measures of functional connectivity: Relevance for studies of neurodevelopment in youth. NeuroImage. 60: 623–632.

66. Ciric R, Wolf DH, Power JD, Roalf DR, Baum G, Ruparel K, et al. (2017): Benchmarking of participant-level confound regression strategies for the control of motion artifact in studies of functional connectivity. NeuroImage. 1–22.

67. Pruim RHR, Mennes M, van Rooij D, Llera A, Buitelaar JK, Beckmann CF (2015): ICA-AROMA: A robust ICA-based strategy for removing motion artifacts from fMRI data. NeuroImage. 112: 267–277.

68. Pruim RHR, Mennes M, Buitelaar JK, Beckmann CF (2015): Evaluation of ICA-AROMA and alternative strategies for motion artifact removal in resting state fMRI. NeuroImage. 112: 278–287.

69. Satterthwaite TD, Elliott MA, Gerraty RT, Ruparel K, Loughead J, Calkins ME, et al. (2013): An improved framework for confound regression and filtering for control of motion artifact in the preprocessing of resting-state functional connectivity data. NeuroImage. 64: 240–256.

70. Hyman SE (2007): Can neuroscience be integrated into the DSM-V? Nat Rev Neurosci. 8: 725–732.

## References

1. Storch EA, Bagner D, Merlo LJ, Shapira NA, Geffken GR, Murphy TK, Goodman WK (2007): Florida obsessive-compulsive inventory: Development, reliability, and validity. J Clin Psychol. 63: 851–859.

2. Ferris JA, Wynne HJ (2001): The Canadian problem gambling index. Ottawa, ON.

3. Hoaglin DC, Iglewicz B (1987): Fine-Tuning Some Resistant Rules for Outlier Labeling. Journal of the American Statistical Association, 2nd ed. 82: 1147–1149.

4. Muthén LK, Muthén BO (2012): Mplus user’s guide, 7 ed. Los Angeles, CA: Muthén & Muthén.

5. Tiego J, Oostermeijer S, Prochazkova L, Parkes L, Dawson A, Youssef G, et al. (2018): Overlapping Dimensional Phenotypes of Impulsivity and Compulsivity. Retrieved from https://osf.io/qa9eh/.

6. Byrne BM (2012): Structural equation modeling with Mplus: Basic concepts, applications, and programming. New York: Routledge.

7. Enders CK (2010): Applied Missing Data Analysis. Guilford Press.

8. Bagozzi RP, Yi Y (2011): Specification, evaluation, and interpretation of structural equation models. J of the Acad Mark Sci. 40: 8–34.

9. Bentler PM, Bonett DG (1980): Significance tests and goodness of fit in the analysis of covariance structures. Psychological Bulletin. 88: 588–606.

10. Browne MW, Cudeck R (1993): Alternative ways of assessing model fit. In: Testing Structural Equation Models. Newbury Park, CA, pp 136–162.

11. Marsh HW, Hau K-T, Wen Z (2004): In Search of Golden Rules: Comment on Hypothesis-Testing Approaches to Setting Cutoff Values for Fit Indexes and Dangers in Overgeneralizing Hu and Bentler’s (1999) Findings. Structural Equation Modeling: A Multidisciplinary Journal. 11: 320–341.

12. Yu CY (2002): Evaluating cutoff criteria of model fit indices for latent variable models with binary and continuous outcomes. University of California Los Angeles.

13. Grice JW (2001): Computing and evaluating factor scores. Psychological Methods. 6: 430–450.

14. Hancock GR, Mueller RO (2001): Rethinking construct reliability within latent variable systems. In: Structural equation modeling Present and future - A festchrift in honor of Karl Joreskog. Lincolnwood, IL, pp 195–216.

15. DiStefano C, Zhu M, Mîndrilă D (2009): Understanding and Using Factor Scores: Considerations for the Applied Researcher. Practical Assessment, Research Evaluation. 14: 1–11.

16. Avants B, Epstein C, Grossman M, Gee J (2008): Symmetric diffeomorphic image registration with cross-correlation: Evaluating automated labeling of elderly and neurodegenerative brain. Medical Image Analysis. 12: 26–41.

17. Parkes L, Fulcher B, Yücel M, Fornito A (2018): An evaluation of the efficacy, reliability, and sensitivity of motion correction strategies for resting-state functional MRI. NeuroImage. 171: 415–436.

18. Power JD, Plitt M, Kundu P, Bandettini PA, Martin A (2017): Temporal interpolation alters motion in fMRI scans: Magnitudes and consequences for artifact detection. PLoS ONE. 12: e0182939–20.

19. Pruim RHR, Mennes M, van Rooij D, Llera A, Buitelaar JK, Beckmann CF (2015): ICA-AROMA: A robust ICA-based strategy for removing motion artifacts from fMRI data. NeuroImage. 112: 267–277.

20. Pruim RHR, Mennes M, Buitelaar JK, Beckmann CF (2015): Evaluation of ICA-AROMA and alternative strategies for motion artifact removal in resting state fMRI. NeuroImage. 112: 278–287.

21. Heim S, Eickhoff SB, Ischebeck AK, Friederici AD, Stephan KE, Amunts K (2007): Effective connectivity of the left BA 44, BA 45, and inferior temporal gyrus during lexical and phonological decisions identified with DCM. Hum Brain Mapp. 30: 392–402.

22. Parkes L, Fulcher BD, Yücel M, Fornito A (2017): Transcriptional signatures of connectomic subregions of the human striatum. Genes, Brain and Behavior. 25: 1176–17.

23. Rodriguez A, Reise SP, Haviland MG (2016): Evaluating bifactor models: Calculating and interpreting statistical indices. Psychological Methods. 21: 137–150.

24. Tabachnick BG, Fidell LS (2013): Using multivariate statistics, 6 ed. Boston: Pearson Education, Inc.

25. DeCarlo LT (1997): On the meaning and use of kurtosis. Psychological Methods. 2: 292–307.

26. Small NJH (1980): Marginal Skewness and Kurtosis in Testing Multivariate Normality. Applied Statistics. 29: 85.

27. Satterthwaite TD, Wolf DH, Loughead J, Ruparel K, Elliott MA, Hakonarson H, et al. (2012): Impact of in-scanner head motion on multiple measures of functional connectivity: Relevance for studies of neurodevelopment in youth. NeuroImage. 60: 623–632.

28. Satterthwaite TD, Elliott MA, Gerraty RT, Ruparel K, Loughead J, Calkins ME, et al. (2013): An improved framework for confound regression and filtering for control of motion artifact in the preprocessing of resting-state functional connectivity data. NeuroImage. 64: 240–256.

29. Ciric R, Wolf DH, Power JD, Roalf DR, Baum G, Ruparel K, et al. (2017): Benchmarking of participant-level confound regression strategies for the control of motion artifact in studies of functional connectivity. NeuroImage. 1–22.

